# Relationship Between Inter-individual Variation in Circadian Rhythm and Sociality: A case Study Using Halictid Bees

**DOI:** 10.1101/2021.09.03.458748

**Authors:** Sofía Meléndez Cartagena, Carlos A. Ortiz-Alvarado, Patricia Ordoñez, Claudia S. Cordero-Martínez, Alexandria F. Ambrose, Luis A Roman Lizasoain, Milexis A Santos Vega, Andrea V Velez Velez, Jenny P. Acevedo-Gonzalez, Jason Gibbs, Theodora Petanidou, Thomas Tscheulin, John T. Barthell, Victor H. González, Tugrul Giray, José L. Agosto-Rivera

**Author notes:** **Correspondence:** Corresponding author: Sofía Meléndez Cartagena.

## Abstract

The bee family Halictidae is considered to be an optimal model for the study of social evolution due to its remarkable range of social behaviors. Past studies in circadian rhythms suggest that social species may express more diversity in circadian behaviors than solitary species. However, these previous studies did not make appropriate taxonomic comparisons. To further explore the link between circadian rhythms and sociality, we examine four halictid species with different degrees of sociality, three social species of *Lasioglossum*, one from Greece and two from Puerto Rico, and a solitary species of *Systropha* from Greece. Based on our previous observations, we hypothesized that species with greater degree of sociality will show greater inter-individual variation in circadian rhythms than solitary species. We observed distinct differences in their circadian behavior that parallel differences across sociality, where the most social species expressed the highest inter-individual variation. We predict that circadian rhythm differences will be informative of sociality across organisms.

## Introduction

Understanding the evolutionary link between solitary and eusocial lineages, and their adaptive behaviors, such as those expressed in reproduction and brood care, is a perennial question in insect evolutionary biology (Toth and Rehan 2017). Insects, and in particular hymenopterans, have been useful in observing how sociality is related to other types of behaviors, such as competitor effects (Peters et al. 2017). A potential behavior to be evaluated in relation to sociality is circadian rhythm, as it has been proposed to be governed by demands arising from sociality not only in insects but also in mammals (Mistlberger 2004; Giannoni-Guzman et al. 2014; Beer and Helfrich-Förster 2020).

Circadian rhythms can be viewed as a biological clock that most living organisms possess. These biological clocks regulate processes such as gene expression, behavior, body temperature, and sleep-wake patterns. Biological clocks follow a rhythm that is approximately synchronized to Earth’s 24-hour rotation using signals from the environment called “zeitgebers” or time givers. This process of synchronization needs active reestablishment, and it is called entrainment (Roenneberg et al. 2003).

Circadian rhythms have been studied in a wide range of organisms, from plants, invertebrates, birds to mammals (Helm and Visser 2010). The traditional model animal to study this phenomenon is the fruit fly, *Drosophila melanogaster*, where biological clocks are described at the molecular level (Dubowy and Sehgal 2017). Although this model has been pivotal to the understanding of circadian rhythms, the lack of genetic diversity in the fruit fly reduces the relevance of the model because it limits questions regarding individual differences in rhythms.

Past studies, such as those of Bloch et al. (2001) and Moore et al. (1998), revealed that the rhythmicity of honey bees changed with age. Additionally, Giannoni-Guzman et al. (2020) showed that foragers in the wild display discrete categories that suggest temporal shift work. An earlier study from Giannoni-Guzman and colleagues (2014) compared the endogenous period of three different variants of honey bees (*Apis mellifera carnica*, *Apis mellifera caucasica* and *Apis mellifera gAHB*) as well as similarly sized insects from different orders and families. They found that honey bees and paper wasps (*Polistes crinitus* and *Mischocyttarus phthisicus*) had a larger degree of circadian period variation within the population in comparison to *D. melanogaster*. The authors mentioned several possible explanations for their observations, one of them being demands of sociality. In a more recent study by Beer and Helfrich-Förster (2020), they explore this connection further and note that the development of the circadian circuitry varies between a eusocial (*Apis mellifera*) and a solitary species (*Osmia bicornis*). In particular, they observe that eusocial individuals do not display circadian locomotion at adult emergence, while the solitary individuals emerge displaying a fully rhythmic circadian rhythm; These differences are attributed to their opposite levels of sociality. However, because these two past studies were done with species spanning from different taxonomic groups, it would be difficult to support their claims without taking phylogeny into account. Nevertheless, these works do give a basis to ask how circadian rhythms vary and are an integral part of the survival strategy and organization of these animals. Moreover, it makes us consider that the level of sociality in different organisms may play a role in their daily activity patterns. An important emerging feature is that in social insects, be it groups of individuals as defined by their age and job (Moore et al. 1998, Bloch et al. 2001), or individuals in the same age and task group (Giannoni-Guzman et al. 2014, 2020), exhibit differences in their circadian rhythms with functional significance for their sociality. We hypothesize that with increasing levels of social organization, as can be measured in size of the social group, greater levels of individual variation in circadian rhythms will be observed.

We tested this hypothesis of socially increased individual variation, by examining circadian rhythms of foragers from differently social halictid bees. Halictidae (Hymenoptera) is a bee family considered to be a great model for the study of social evolution due to its exceptional diversity in respect to social behavior within and among species and populations (Schwarz et al. 2007). *Lasioglossum* Curtis is one of the two genera in the tribe Halictini that displays eusocial behavior, but also includes solitary representatives and a range of intermediate social categories (Danforth et al. 2003; Gibbs et al. 2012). Additionally, past studies have shown plasticity in the social behavior even among populations of the same species (Eickwort et al. 1996; Field 1996; Field et al. 2010; Richards et al. 2003; Soucy and Danforth 2002; Richards 2000). Depending on environmental conditions, such as elevation, latitude and seasonality, halictid bees might display different modes of sociality. Species with social nests may revert to solitary behavior at high latitudes and altitudes (Eickwort et al. 1996; Packer et al. 1983; Field et al. 2010) or based on access to mates (Yanega 1988/1989). Jeanson et al. (2008) studied members of a solitary species, *Lasioglossum* (*Ctenonomia*) sp. NDA-1 and observed the results of having them nest in pairs. They finally observed that after some time together, the individuals in the nest started to show signs of division of labor. This plasticity and diversity of behavior in addition to the close taxonomic relation, makes Halictidae an optimal model for observing the relationship between sociality and circadian rhythm (Bloch and Grozinger 2011).

To better understand association of sociality with individual variation in circadian rhythms, we have set out to document the rhythm in four species of halictid bees that span a gradient of social complexity. *Systropha curvicornis* (Scopoli) (Halictidae: Rophitinae), a solitary pollinator specialist (Grozdanić and Mučalica 1966) considered ancestrally solitary within the family Halictidae (Patiny and Michez 2007; Patiny et al. 2008; Danforth et al. 2008), and three species of *Lasioglossum (L. ferreri, L. enatum* and *L. malachurum*), which were selected because of their varying levels of social behavior (Eickwort 1988; Wyman and Richards 2003; Gibbs 2018). Species of *Lasioglossum* likely had a common ancestor capable of eusocial nesting but have reverted multiple times to other levels of sociality (Danforth et al. 2003; Brady et al. 2006; Gibbs et al. 2012). *Lasioglossum (Dialictus) ferrerii* (Baker) and *L. (Dialictus) enatum* (Gibbs) occurs in the Caribbean whereas *L. malachurum* across Europe; the first species nests communally, that is each individual contributes to nest construction and reproduction (Michener 1974; Eickwort 1988). Although *L. enatum* has not been thoroughly studied, this species is part of a species complex that includes weakly eusocial species. Namely, *L. gemmatum* (Smith) and *L. parvum* (Cresson) from Jamaica and the Bahamas, which exhibit reproductive division of labour (Eickwort 1988). There is no evidence of morphologically defined castes beyond reproductive status in these two species and thus we assumed this is likely the case for *L. enatum* in Puerto Rico, where we conducted our experiments. In contrast, *L. malachurum* (Kirby) is an obligately eusocial species with morphologically well-defined queen and worker castes (Richards 2000; Wyman and Richards 2003). *Lasioglossum malachurum* is known to display varying degrees of behaviors depending on location (Richards 2000). In Lesbos, Greece, where we studied this species, they were observed to exhibit a facultatively eusocial behavior (Wyman and Richards 2003). In summary, the level of sociality in the four species goes from lowest to highest: solitary *Systropha curvicornis*, communal *Lasioglossum ferreri*, weakly eusocial *Lasioglossum enatum*, and the eusocial *Lasioglossum malachurum*. The socially increased individual variation hypothesis predicts individual variation of foragers from each species to increase from solitary to eusocial. Alternatively, individual variation could be the result of ecological differences such as differences in robustness of time givers in different environments. Two of the halictids examined here are from a tropical zone (a communal and an eusocial) and two are from a temperate zone (a solitary and an eusocial).

The social plasticity of *Lasioglossum* and its potential as a model for social evolution leads us to believe that observing this group of bees can give invaluable insight on how social behavior affects biological clocks. To test if sociality increases individual variation, we captured the forager bees as they were visiting flowers and observed their circadian variability in the laboratory using constant conditions.

## Methods and Materials

### Study sites

#### Puerto Rico

*Lasioglossum ferrerii* and *L. enatum* were captured using 15 mL falcon tubes from flowers at the Balneario de Luquillo parking lot (18.38706 N 65.72517W, 3 Meters) in Puerto Rico. This site is characterized by having many vine-type plants, high vegetation density, and it is located right next to a road with *Momordica charantia* being the most abundant (Figure 1.A and 1.B). Most bees were caught between 8:00 and 12:00 h at the flowers of *Momordica charantia* L. (Cucurbitaceae), *Sida acuta* Burm.fil. (Malvaceae), and *Bidens alba* (L.) DC. (Asteraceae). We also observed them visiting *Euphorbia heterophylla* which has not been reported in previous literature. Collections took place during the months of December, January, March, and August. In total we collected 36 bees, 26 of which were *L. ferrerii* and 10 were *L. enatum*.

**Figure 1:**
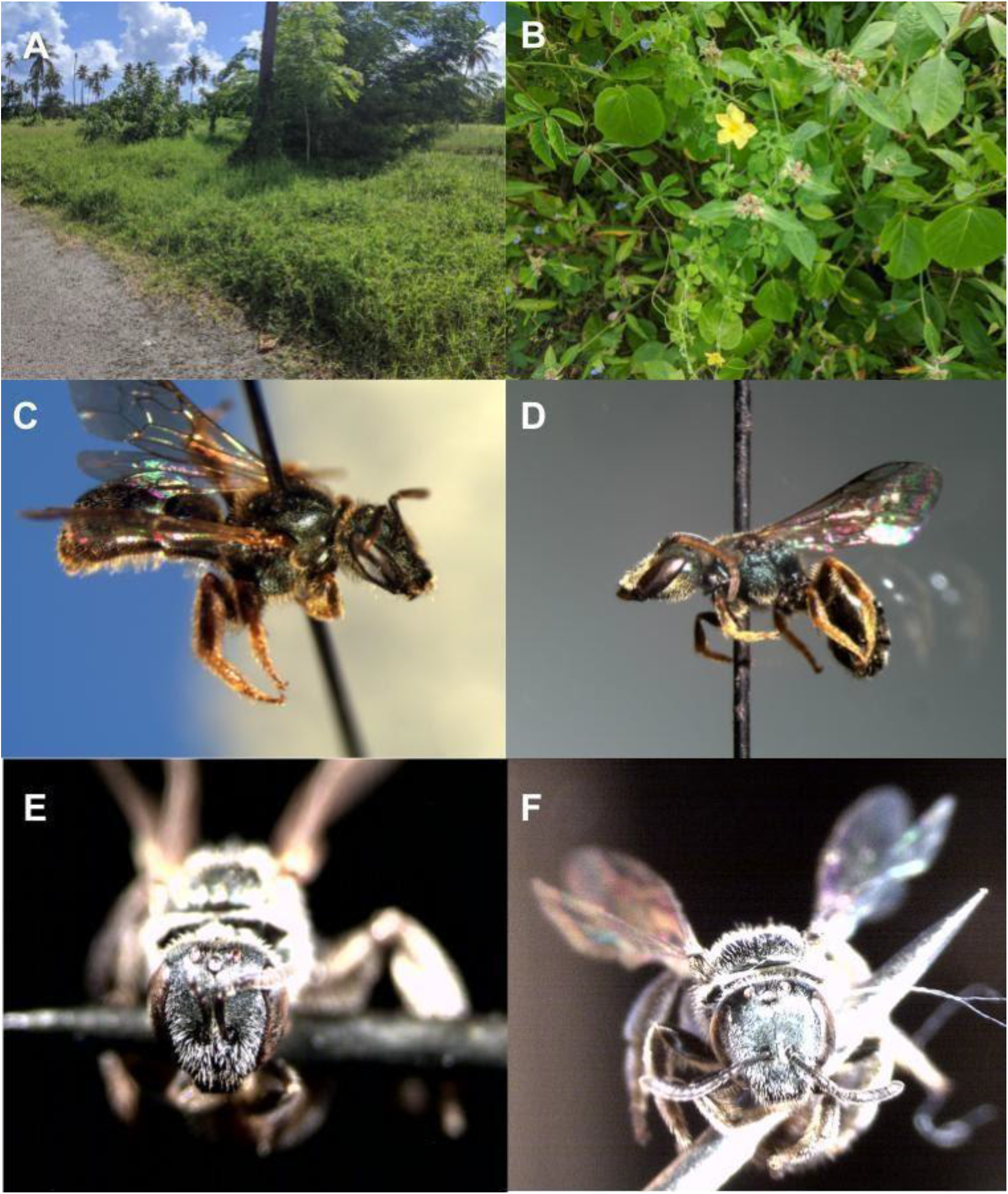
Habitat (A-B) and Species Observed (C-F). **A)** Puerto Rico study site in which *L. ferrerii* and *L. enatum* were captured. **B)** Some of the vegetation the bees were observed visiting, with flowers belonging to the families: Commelinaceae, Cucurbitaceae and Euphorbiaceae. **C)** Female of *L. ferrerii* distinguished from the male by its short antenna and pointed abdomen. **D)** Male of *L. ferrerii*, distinguished by its long antennae and flat abdomen. This species is known for its long head shape and metallic metasoma (Gibbs 2018). **E)** Female of *L. enatum* distinguished from the male by its short antenna and pointed abdomen. **F)** Male of *L. enatum*, distinguished by its long antennae and flat abdomen. This species is distinguished by: “tegula punctate, extended posteriorly to form a small angle, mesepisternum punctate and metasoma brown” (Gibbs 2018).

#### Greece

*Systropha curvicornis* and *L. malachurum* bees were collected between 6:00 and 9:00 h from flowers of *Convolvulus arvensis* (Convolvulaceae) that were growing on a recently cut wheat field in Skala Kallonis (39° 10’N 26° 20’E, 0 Meters) on the Island of Lesbos, Greece. We used 15 mL falcon tubes to catch bees in the field, which would house the individual for the duration of the experiment. Sampling was conducted on July 3 of 2017. From this sampling 118 bees were *L. malachurum* and 34 were *S. curvicornis*.

### Laboratory settings

After collection in the falcon tubes, bees were provided with food that lasted for the whole of the observation period. The food recipe we used varied between the studies in Puerto Rico and Greece (As explained below). The main nutrient for both recipes was sugar and therefore are nutritionally comparable. However the agarose based recipe (Puerto Rico) was more convenient in terms of ease and speed of preparation due to the fact that an independant water system was not necessary.

Food preparation varied by locations as follows: In Puerto Rico, for every 0.89 ml of water, 1 g of sucrose and 0.1 g of agarose were used. The water was heated in a stirring plate with a magnetic stirrer placed at the bottom. We added the sucrose first to the solution, and when it was dissolved, the agarose was incorporated. The solution was left stirring until it turned into a lighter color while mindful of not letting the solution heat too much, or part of the volume would be lost. As a form of assurance, we made 3 ml more than what was expected to be used. After all solids were diluted, we quickly pipetted 1 ml into the bottom of a 15 ml centrifuge tube, being mindful of not letting it splash. Once all of the tubes had their portion of the solution, they were allowed to reach room temperature and finally they were refrigerated. The final product was a gel that could be kept refrigerated until the day it needed to be used as long as it did not dehydrate.

In Greece, captured *S. curvicornis* and *L. malachurum* were fed with ApiYem (Namik Kemal University with Kosgeb R&D Innovation Project) which is a commercial bee feed composed of 78.5% sugar and 21.5% invert syrup. Food was placed in the cap-end of each tube and a damp cotton was placed in the other end of the tube to provide water to the bees. The water supply was refilled every 2–3 days. Resources were provided ad libitum during the complete running period of the experiment.

#### Locomotor activity monitoring

Each bee was monitored individually for at least seven days in the falcon tube in which they were captured. The tubes were plugged using cotton balls to let air circulate. These tubes were then placed into TriKinetics’ Locomotion Activity monitors (LAM16) that, in turn, were put inside incubators that were set to constant conditions. In Greece the temperature was 26, humidity 78%, constant darkness. In Puerto Rico the temperature was 30, humidity 65%, constant darkness. The differences in the environmental chamber conditions were set to resemble the average daytime parameters at each location.

#### Species Identification

The individuals caught in Puerto Rico were identified using Gibbs (2018). Samples collected in Greece were identified by an in-field expert, Victor H. Gonzalez (University of Kansas).

### Data processing and analysis

#### Circadian Analysis

Circadian rhythm and locomotor activity for our subjects were analyzed using the MATLAB toolboxes developed in Jeffrey Hall’s laboratory (Levine et al. 2002). The outputs provided data on the individual’s locomotor activity throughout the experiment in the form of an actogram, average activity plot, and an autocorrelation that also calculates rhythm strength.

To test if the observed differences in circadian patterns across species were statistically significant, we applied a Brown-Forsythe one way ANOVA with a Dunnett’s T3 multiple comparisons test using GraphPad Prism version 8.4.3 for Windows, GraphPad Software, San Diego, California, USA, www.graphpad.com. The variables taken into consideration for this was time, species, individuals, and interspecies variation.

## Results

During the Summer of 2017 on the 3^rd^ of July between the hours of 6:00 am and 9:00 am, *S. curvicornis* and *L. malachurum* were caught as they visited the flowers of Convolvulus *arvensis o*n the island of Lesbos. They were transported from the field to the laboratory and placed inside an incubator for 10 days of which 8 were in constant conditions (26 °C and 57 % humidity) with the purpose of characterizing their intrinsic biological clockDuring this collection 118 bees were *L. malachurum* although only 98 survived until the end and 34 were *S. curvicornis* of which only 4 females survived the study period.

Figure 2.A illustrates the average of individuals evaluated under constant conditions of the Greek, solitary and specialist pollinator *S. curvicornis*. Its period runs slightly short at approximately 22 hours, with the peak of its activity in the early morning and an average rhythm strength of 4.4. The population average is consistent with the individuals examined, as illustrated in Figure 2.B; a randomly selected individual looks fairly similar to the activity plotted for the population average. The average for period was 22.75 with a standard deviation of 0.41, Rhythm Strength had an average of 4 and standard deviation of 0.707.

**Figure 2:**
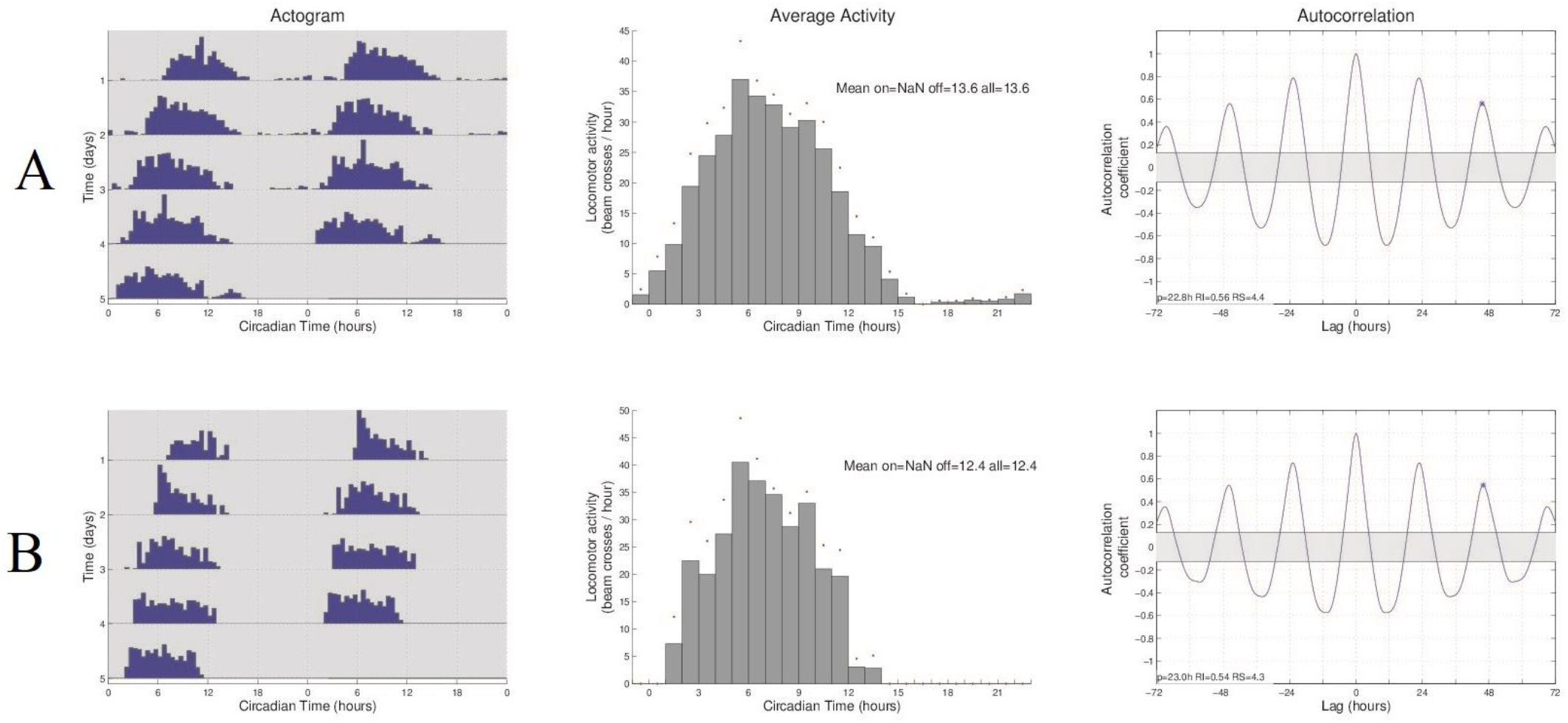
Female *S. curvicornis* exhibit short period phenotype under constant darkness (<24 h endogenous circadian rhythm). (i) Double-plotted actogram showing the locomotor activity pattern of: (A) The average of all 4 individuals examined of *S. curvicornis*. (B) A representative individual randomly selected from the population. In a double plotted actogram, each row represents locomotor activity (counts per 30 min) of two consecutive days and the second is repeated such that it is always the beginning of the next row. The x-axis shows the time of day under constant darkness expressed as circadian time (CT). (ii) Average of the locomotor activity patterns of the five days observed into one. The y-axis represents activity over time and the x-axis represents CT (iii) Autocorrelation plots used to determine the period (p), rhythm index (RI) and rhythm strength (RS). The oscillations indicate periodicity. The asterisk at the third peak of the autocorrelation plot indicates the particular time point used for the determination of the rhythm parameters.

The consistency displayed by our population of female *S. curvicornis* is contrasted with the diversity observed in the other 3 species analyzed in this study. This is especially true for the eusocial *L*. *malachurum*, for whom after careful evaluation of the data we had to create a classification schematic (Figure 3) to appropriately describe the phenotypes being displayed by the population. The population average shows that *L. malachurum*, as a species, has a perfectly circadian 24-hour period under constant dark conditions. Peak average activity of *L. malachurum* is at 6:00 h, with no clear rest periods when all individuals are averaged. When examined individually, we found five distinct patterns of circadian activity patterns (Figure 3.A and Figure 4). These patterns can be divided into 2 large branches (Figure 3.A), those that are rhythmic and those that are arhythmic i.e, individuals with uniform distribution of activity. Rhythmic individuals varied in the amplitude of their activity rhythm and therefore were classified as strong or weakly rhythmic. Moreover, both strong and weak categories are subdivided into unimodal or bimodal based on the number of activity peaks per day. For example, a bimodal individual is active during two different instances of the day like the morning and the afternoon (Figures 4.B and 4.C), while a unimodal individual is mostly active during a set time of the day (Figures 4.D and 4.E).

**Figure 3:**
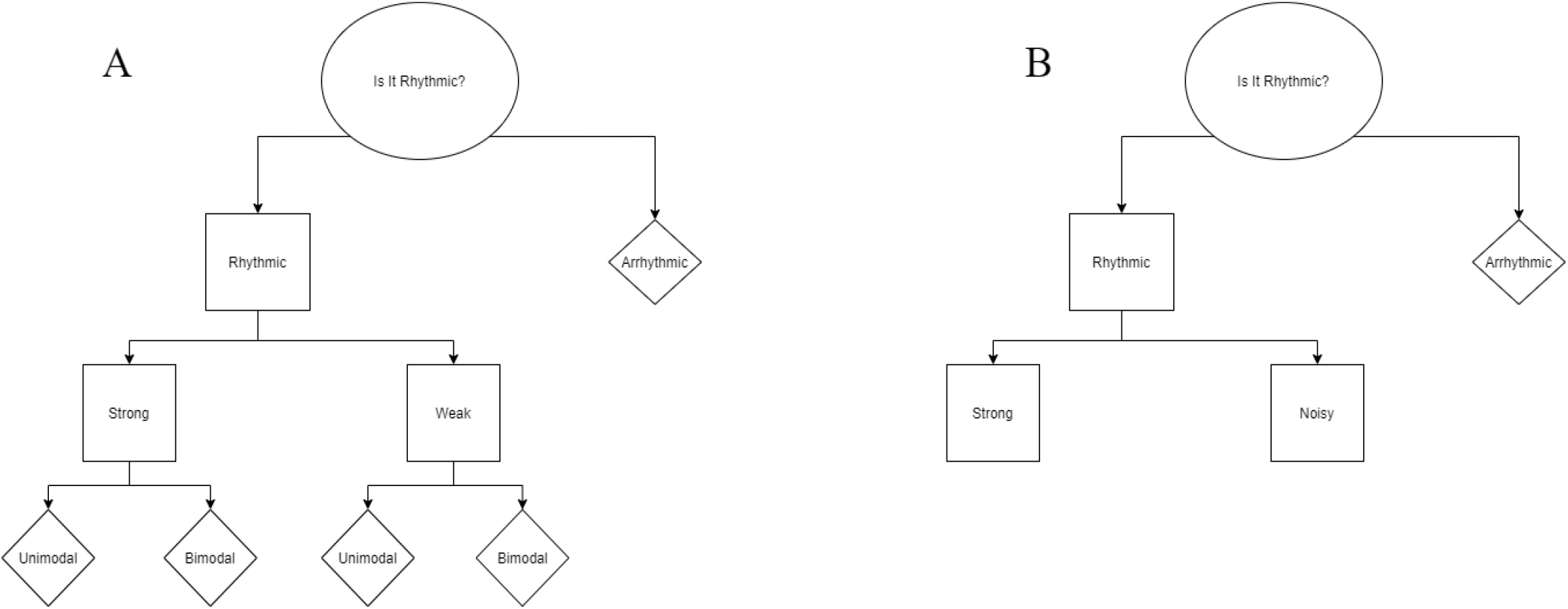
A summary of the variations in the circadian rhythm as observed in: **A)** *Lasioglossum malachurum*, **B)** *Lasioglossum enatum* and *Lasioglossum ferreirii*. The circle represents the root of the flowchart, squares represent nodes that branch off and rhombuses represent leaves. In total for malachurum, 5 distinct behaviors were observed.

**Figure 4:**
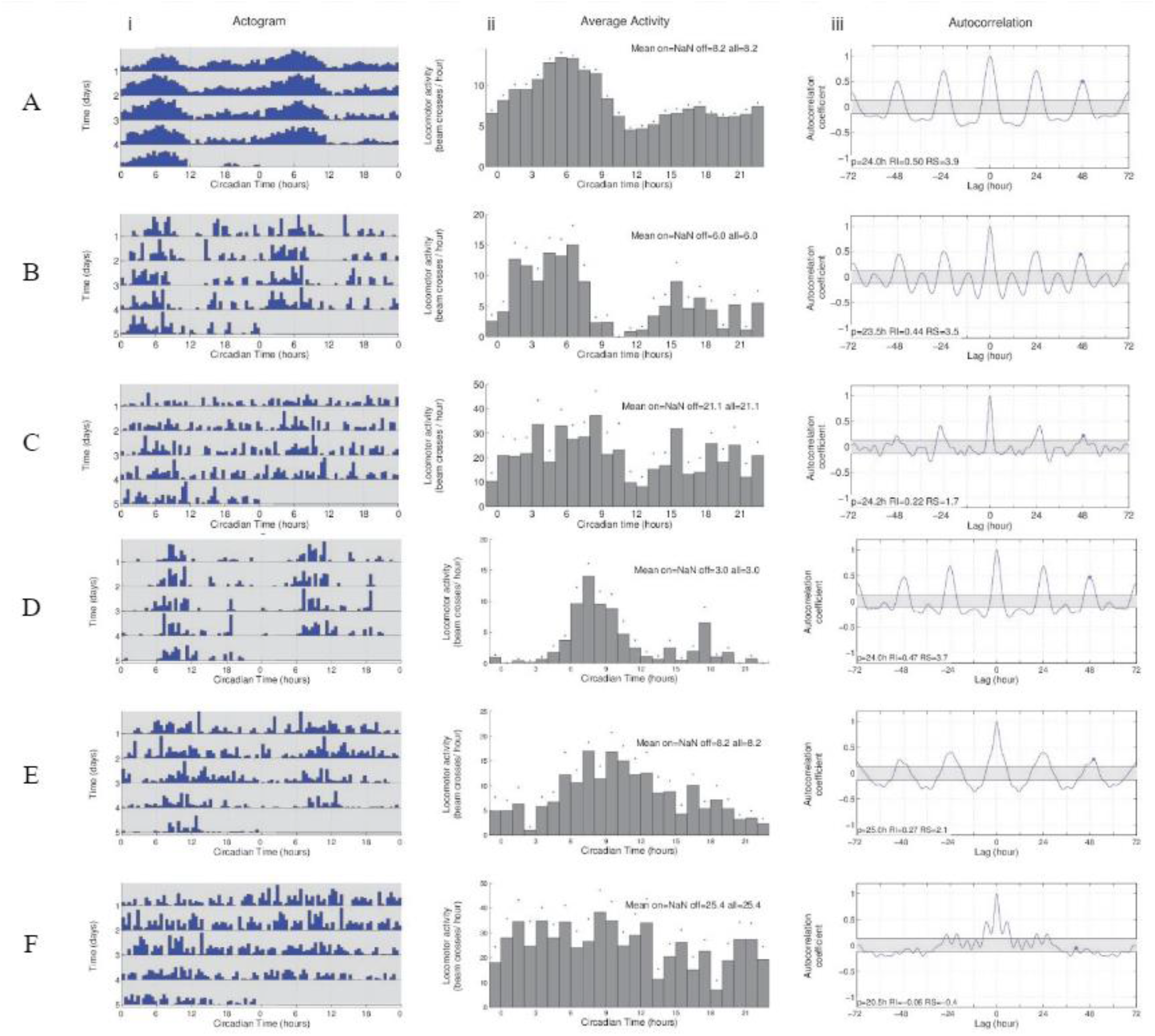
*L. malachurum* exhibits a variety of circadian phenotypes under constant dark conditions. (i) Double-plotted actogram showing the locomotor activity pattern for 5 days of: (A) The average of all 98 individuals examined and representatives for the following categories: (B) Bimodal Rhythmic (C)Weakly Rhythmic Bimodal, (D) Unimodal Rhythmic, (E) Weakly Rhythmic Unimodal, (F) and Arrhythmic circadian behaviors. (ii) An average activity plot for the five days of observation (iii) Autocorrelation plots used to determine the period (p), rhythm index (RI) and rhythm strength (RS).

Strongly rhythmic individuals (either unimodal or bimodal) constituted 41% of individuals. These patterns are recognized by a strong Rhythm Strength (RS)(Figures 4.B.iii and 4.D.iii) on average higher than 2.67 and clear rest and active periods both in the double plotted actogram (Figures 4.B.i and 4.D.i) and the average activity plot(Figures 4.B.ii and 4.D.ii). Weak rhythmicity (Figure 4.C and 4.E) was observed in 21.6% of individuals and they were characterized by having RS values (Figure 4.C.iii and 4.E.iii) that average on 1.79, but their actograms (Figure 4.C.i and 4.E.i) do not show a clear pattern of locomotor activity. Finally, 38% of individuals were arrhythmic. Both the double plotted actogram (Figure 4.F.i) and the average activity plot (Figure 4.F.ii)for arrhythmic bees do not have any discernible daily pattern of activity or inactivity. Often the autocorrelation (Figure 4.F.iii) does not return any values.

On February 19, 2020, between 8:00 am and 10:00 am, at the Balneario de Luquillo (Figure 1.A and B), 36 bees were captured as they visited *Bidens alba*, *Momordica charantia* and *Sida acuta*. Bees were captured and monitored individually in one tube each (modified from Giannoni-Guzman et al. 2014). In the laboratory, the bees were placed inside an incubator for seven days in constant conditions so we could characterize their intrinsic biological clock.

Of the 36 bees captured, 26 were identified as *Lasioglossum ferrerii* (Figure 1.C and 1.D) and ten as *Lasioglossum enatum* (Figure 1.E and 1.F). Only 22 *L. ferrerii* and 8 *L. enatum*, survived the entire observation period and were used for analysis. The average peak of circadian activity for *L. ferrerii* is between 6:00–7:00 (Figure 5.A.ii) with a 23 hour period (Figure 5.A. iii), making it short. Individuals fell into two categories, 50% were strongly rhythmic and 50% were weakly rhythmic (Figures. 5B and 5C). The peak of average activity for *L. enatum* is from the fifth to the seventh hours of the day, with a circadian period of 23.8 hours (Figure 6.Aiii), just slightly short of a day. The average peak of activity for *L. enatum* was 6:00-7:00 (Figure 6.A.ii). *L. enatum* also had three patterns of activity with similar characteristics to that of the umbrella categories for *L. malachurum*, and we saw it fit to categorize them in a similar fashion (Figure 6). 12.5% of the observed population fell into the strongly rhythmic category, 25% in the weak rhythmic category and 62.5% in the arrhythmic category.

**Figure 5:**
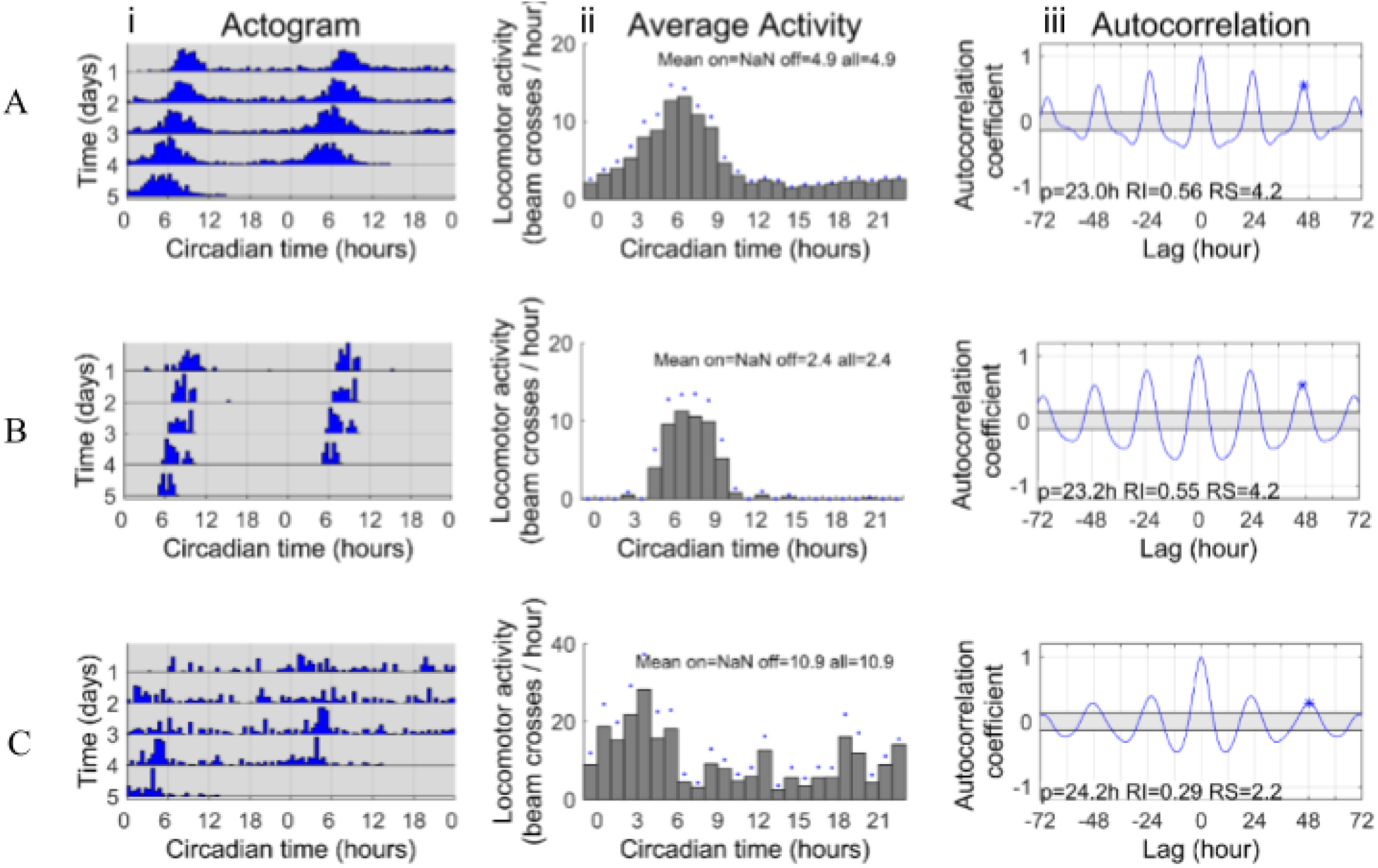
Description of the circadian behaviors exhibited by *L. ferrerii* under constant dark conditions. (i) Double-plotted actogram of the locomotor activity from the five-day observational period for: **A)** All 22 individuals from the data set averaged out into one representative individual. **B)** A representative individual out of the 11 from the category Strongly Rhythmic. **C)** A representative individual out of the 11 from the category Noisy Rhythmic. (ii) An average activity plot for the five days of observation (iii) Autocorrelation plots used to determine the period (p), rhythm index (RI) and rhythm strength (RS).

**Figure 6:**
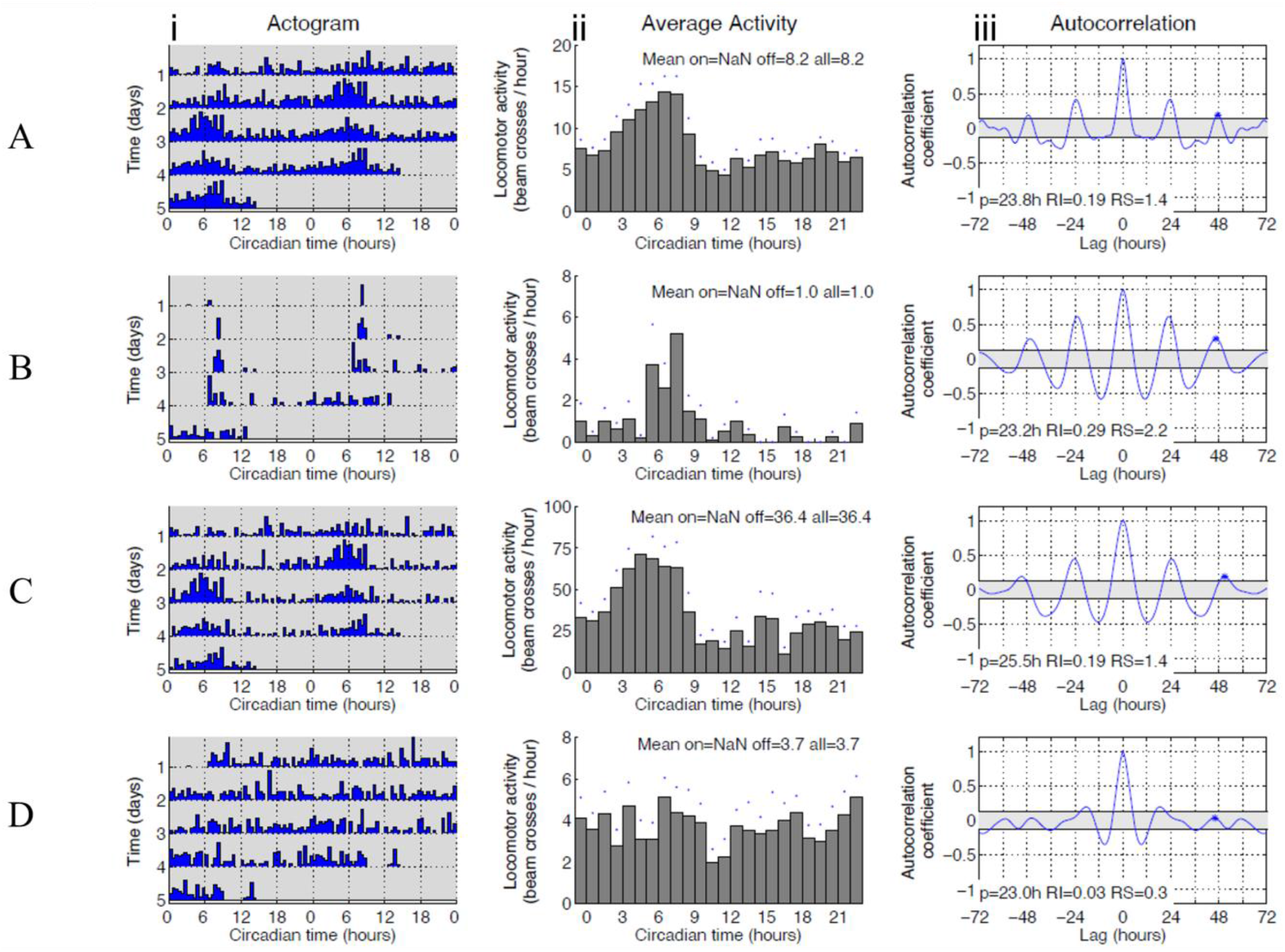
Description of the circadian behaviors exhibited by *L. enatum* under constant dark conditions. (i) Double-plotted actogram of the locomotor activity from the five-day observational period for: **A)** All 8 individuals from the data set averaged out into one representative individual. **B)** The only individual from the category Strongly Rhythmic. **C)** A representative individual out of the 2 from the category Noisy Rhythmic. **D)** A representative individual out of the 5 from the Arrhythmic category. (ii)An average activity plot for the five days of observation (iii) Autocorrelation plots used to determine the period (p), rhythm index (RI) and rhythm strength (RS).

In summary, all four of the described species followed unique patterns of behavior (Figure 7.A) characterized by the amount of interindividual variation. *Systropha curvicornis* was the species with the least amount of observed interindividual variation in its daily activity patterns, followed by *L. ferrerii* with two distinct patterns of behavior, then *L*. *enatum* with 3 and lastly *L. malachurum* with 5.

**Figure 7:**
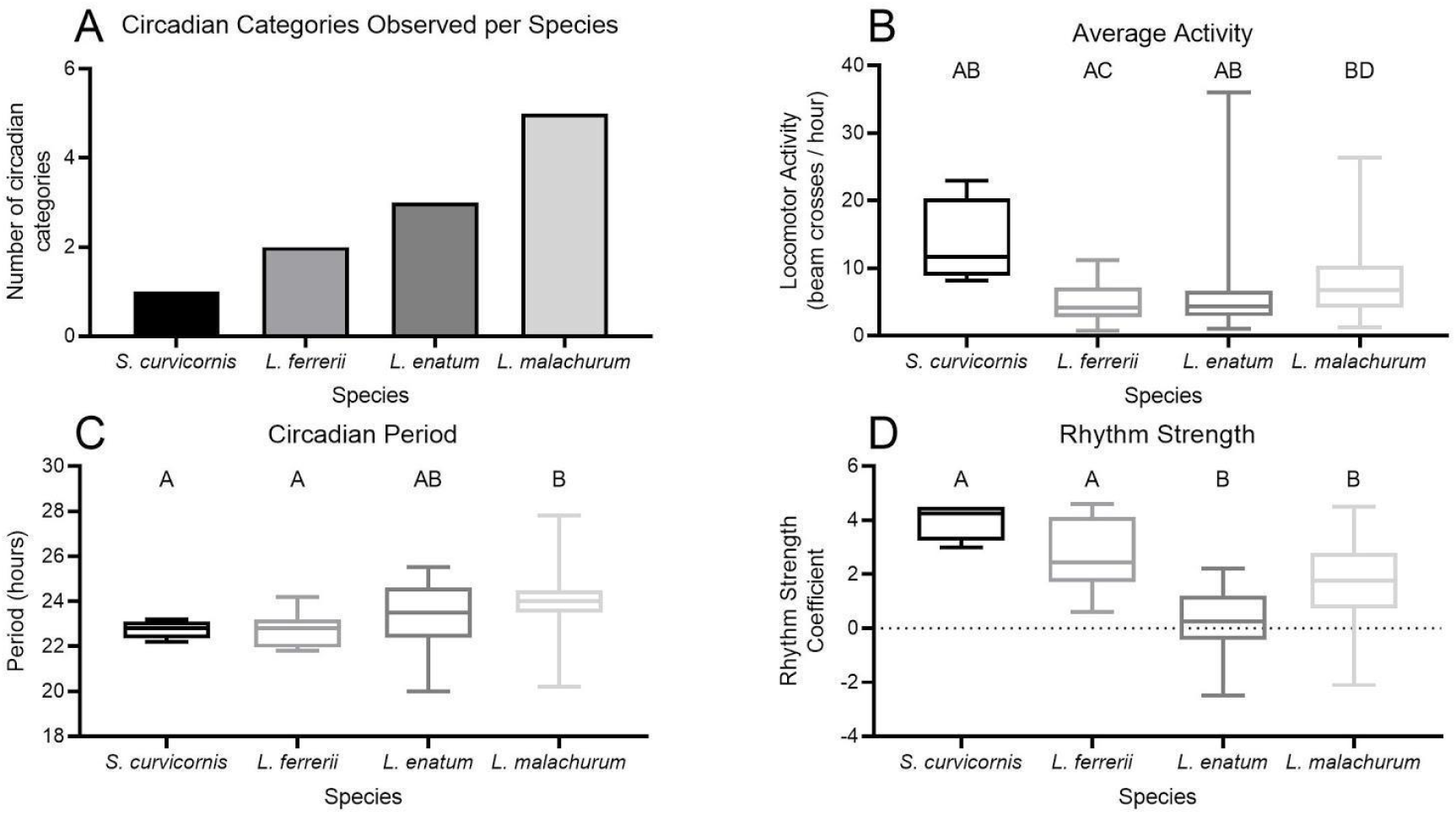
Summary of descriptive and inferential statistics. **A)** Number of circadian categories observed by species. **B)** Box plot illustrating the difference in average locomotor activity between species. *S. curvicornis* has a minimum of 8.200, 25% percentile of 8.900, mean of 13.63, 75% percentile of 20.28 and a maximum of 22.90. *L. ferrerii* has a minimum of 0.7000, 25% percentile of 2.675, mean of 4.950, 75% percentile of 7.100 and a maximum of 11.20. *L.enatum* has a minimum of 1.000, 25% percentile of 2.875, mean of 8.113, 75% percentile of 6.650 and a maximum of 36.00. *L, malachurum* has a minimum of 6.700, 25% percentile of 10.35, mean of 8.201, 75% percentile of 10.35 and a maximum of 26.30. There was only a statistical difference between *L. ferrerii* and *L. malachurum* with a p-value of 0.0016, DF of 61.66 and t of 3.867. **C)** Box plot illustrating the difference in circadian period between species. *S.curvicornis* has a minimum of 22.20, 25% percentile of 22.35, mean of 22.75, 75% percentile of 23.10 and a maximum of 23.20. *L. ferrerii* has a minimum of 21.80, 25% percentile of 21.95, mean of 22.69, 75% percentile of 23.20 and a maximum of 24.200. *L.enatum* has a minimum of 20.00, 25% percentile of 22.40, mean of 23.31, 75% percentile of 24.60 and a maximum of 25.50. *L, malachurum* has a minimum of 20.20, 25% percentile of 23.50, mean of 24.00, 75% percentile of 24.50 and a maximum of 27.80. Both S. *curvicornis* and *L. ferrerii* were significantly different from *L. malachurum* with p-values of; 0.0102 and <0.0001, DFs of; 5.652 and 52.94 and, t of; 5.179 and 6.565, respectively. **D)** Box plot illustrating rhythm strength among species. *S. curvicornis* has a minimum of 3.000, 25% percentile of 3.250, mean of 4.000, 75% percentile of 4.500 and a maximum of 4.500. *L. ferrerii* has a minimum of 0.6000, 25% percentile of 1.700, mean of 2.691, 75% percentile of 4.125 and a maximum of 4.600. *L. enatum* has a minimum of −2.500, 25% percentile −0.4250, mean of 0.2125, 75% percentile of 1.200 and a maximum of 2.200. *L, malachurum* has a minimum of −2.100, 25% percentile of 0.7250, mean of 1.754, 75% percentile of 2.800 and a maximum of 4.500. The solitary, *S. curivcornis*, and communal *L. ferrerii*, were significantly different from the eusocial species, but not each other. Likewise, *L. enatum* and *L. malachurum* were not significantly different.*S. curvicornis* vs. *L. enatum*; p-value of 0.0014, df of 7.932 and t of 6.226. *S. curvicornis* vs. *L. malachurum*; p-value of 0.0177, df 4.04 of and t of 5.895. *L. ferrerii* vs. *L. enatum*; p-value of 0.0056, df of 17.09 and t of 3.982. *L. ferrerii* vs. *L. malachurum*; p-value of 0.0259, df of 33.42 and t of 3.054.

### Cross species comparison of observed circadian parameters

Average activity was only significant between *L. ferrerii* and *L.malachurum* with a p-value of 0.0016 (Figure 7.B). Circadian Period (Figure 7.C) on the other hand showed differences between *S. curvicornis* and *L. ferrerii* with a p-value of 0.0102 as well as *L. malachurum* and *L. ferrerii* with a p-value of <0.0001. Lastly, with even more differences still, Rhythm Strength (Figure 7.D) presented differences between S. *curvicornis* and *L. enatum* (p-value = 0.0014), *S. curvicornis* and *L. malachurum* (p-value = 0.0177), *L. ferrerii* and *L. enatum* (p-value = 0.0056) and finally, *L. ferrerii* and *L. malachurum* (p-value = 0.0259).

## Discussion

This study lays the groundwork for the use of halictid bees for cross species comparisons of circadian rhythms. Consistent with our hypothesis, we found that greater degrees of sociality are associated with larger individual differences of circadian rhythms within a population.

The solitary specialist, *S. curvicornis* as observed in this work, suggests that at least for the females, the population is consistent, displaying a single circadian activity phenotype (Figure 7.A). Activity of these bees is highly rhythmic and shows little variation across samples with an average RS of 4.4 and the peak of activity appears to be near the hour 6 of the day. Overall, the species exhibits a short period phenotype under constant darkness. This high degree of rhythmicity might be due to *S. curvicornis’* evolutionary history as a foraging specialist of *C. arvensis*, which blooms for a brief period during the morning, a pattern described for another closely related species (*S. planidens*: Gonzalez et al. 2014). A rigorous internal clock is important to be able to anticipate the time when resources are available. For example, the immediate development of *Osmia bicornis’* circadian rhythm (Beer and Helfrich-Förster 2020) may be related to nurishment accessibility, as it has been shown in the past that large quantities of pollen are the key to proper larvae development rather than diversity of pollen (Radmacher and Strohm 2010). All three of these species (*S. curvicornis, S. planidens* and *O. bicornis*) lead solitary lifestyles and consequently must assure the survival of their progeny in an individual manner. A strong circadian rhythm can ensure that a female may find sufficient resources efficiently to feed its young.

All three species of *Lasioglossum* examined were shown to have more than one distinct pattern of circadian activity. The most diverse of the three was the facultatively eusocial *L. malachurum* (Figure 7.A) with 5 distinct circadian behaviors. Two of the sub-categories of these behaviors fall under the strongly rhythmic category, which we are calling binomial and unimodal. These rhythmic categories are characterized for having an easy to distinguish pattern in the actogram, clear rest/activity periods in the average activity plot, and an RS higher than one. Another two of the sub-categories fall under the weakly rhythmic umbrella. This umbrella, just like the strongly rhythmic category, can be divided into bimodal and unimodal. These categories can be identified by an actogram with no clear pattern, an average activity plot with more or less clear rest/activity pattern, and an RS larger than one. Lastly, there is the arrhythmic category where no discernible pattern can be pinpointed in the actogram nor in the average activity plot and its RS is less than one. A conceptual map on how these categories are identified can be found in Figure 3.A.

To make descriptions comparable across species, we used the same metrics to categorize the other two bees examined in this study. In categorizing *L. ferrerii* and *L. enatum*, our categories worked as a good basis. *L. ferrerii* only had two distinguishable patterns: rhythmic and noisy rhythmic (Figure 7.A). We decided to change the name from weakly rhythmic to noisy rhythmic because it is a better descriptor (Figure 3.B). Similarly, *L. enatum*, which lives in the same environment as *L. ferrerii*, has 3 distinguishable categories (Figure 7.A). These categories are rhythmic, noisy, rhythmic and arrhythmic. Contrary to *L. ferrerii*, *L. enatum* expressed 5 individuals in the arrhythmic category. Taking into consideration that both of these species of bees were caught in the same environment, and they belong to the same genus, the results suggest that something other than environmental variables are behind these differences.

The difference in expression of circadian patterns between *L. ferrerii* and *L. enatum* could be explained by competition. Both of these species share the same niche in Luquillo, to the point of them being caught in the same flowers during the same range of time. Having a slight difference in rhythmicity can lower the possibility of temporal competition when foraging. *L. ferrerii* on average would be active from 5:00 am to 10:00 am, while average time of activity for *L. enatum* would be from 3:00 am to 8:00 am. Due to that two-hour disphase, it would appear to be less likely that bees from these two species try to visit a flower simultaneously, yet their schedules still have some overlap. These observations are echoed by another study conducted in Greece where they demonstrated that 3 species of carpenter bees (*Xylocopa* spp.) that share the same resources have different circadian rhythms when measured under natural field conditions and also in artificial constant and oscillating conditions (Ortiz-Alvarado et al. *in rev*.).While the solitary *Xylocopa* species have interspecies variation in their circadian rhythms, two out of the 3 examined in Ortiz-Alvarado et al. follow a similar pattern as S. curvicornis, were there isn’t much if any individual differences in the populations examined. Therefore in that particular case, competitor effects can explain the differences in rhythm across species, but in the case of *L. ferrerii* and *L. enatum* it cannot explain the individual differences observed at the species level.

At a higher level looking at the statistical analysis of all 4 halictid bees (Figure 7), some interesting patterns can be noted. In terms of average activity (Figure 7.B), there was only a difference between *L. ferrerii* and *L. malachurum*, none of the other possible combinations of differences occurred. However, the length of the whiskers in the box plots for both *L. enatum* and *L. malachurum* does suggest a level of diversity at the intraspecies level and could be reflective of the number of circadian behaviors observed in these species.

When analyzing the circadian period, the observed differences were between *L. ferrerii* and *L. malachurum* as well as *S. curvicornis* and *L. malachurum* (Figure 7.C). The latter of these pairs have shared environmental conditions when the former pair does not. It is also interesting to note that *L. ferrerii* and *S. curvicornis* cannot be found in the same locations, and yet they do not appear to have significantly different circadian periods, in fact for the populations examined, they appear to be comparable.

The connection between individual differences in circadian rhythm and sociality becomes clearer still when observing rhythm strength (Figure 7.D). Where those species with a lesser number of circadian phenotypes are more similar to each other and likewise the ones with the most diversity are more similar to each other. In other words, *S. curvicornis* and *L. ferrerii* were both significantly different to *L. enatum* and *L. malachurum*, but not to each other. Likewise, there was no significant difference between *L. enatum* and *L. malachurum*. Because there is this consistency of differences that is not associated with differences in environments, we believe that the key to explaining the difference in diversity of behaviors may not lay in competition, but in something more endogenous of the species. Nevertheless, more data is needed.

*Lasioglossum* as a genus is well-known for having a large diversity in social behaviors. This diversity in sociality may also be reflected in other types of behaviors and could be the key to explaining the individual differences in circadian rhythm we observed across foragers within the species. One caveat for the present work is the sample size and populations evaluated (one population for each species and low number of individuals, particularly for *L. enatum*). One future direction is to identify additional populations of the same 4 species to examine consistency of the presented findings. In future studies we will focus on streamlining the process of describing the diverse circadian behaviors observed in a species to facilitate studies with a higher volume of observations. Additionally, we will continue describing the circadian rhythm of additional species of *Lasioglossum* that present social behaviors not evaluated in this study. Understanding the functional relationship between sociality and rhythm will add to further understanding of mechanisms underlying social organization.

## Acknowledgements

Kristie Shirley, Sarah Markland, Jordan Twombly-Ellis, Elif Kardas, Jonathan A. Lopez Colon, Eddie Perez Claudio, Alejandro Armas, Melanie E. Martinez Arrocho, Luis A Roman Lizasoain, Elvia Meléndez-Ackerman, Genetics Laboratory at the University of Puerto Rico, BIOL 4999-423: Scientific Writing-Fall 2020

## Availability of data and material

All data analyzed during this study is included in its supplementary information files.

## Authors’ contributions

All authors contributed to the study conception and design. Material preparation, data collection and analysis were performed by Sofía Meléndez Cartagena, José L. Agosto-Rivera, Carlos A. Ortiz-Alvarado, Claudia S. Cordero-Martínez, Alexandria F. Ambrose, Luis A Roman Lizasoain, Milexis A Santos Vega, Andrea V Velez Velez. The first draft of the manuscript was written by Sofía Meléndez Cartagena and all authors commented on previous versions of the manuscript. All authors read and approved the final manuscript.

## Significance Statement

Circadian rhythms differ across solitary and social organisms. One important feature we discovered is that in highly eusocial insects there was a high level of individual variation in circadian activity. We took advantage of varying levels of sociality within halictids and examined the circadian rhythms of multiple members of the genus *Lasioglossum* and of a *Systhropha* species. This work is the first report to link interindividual variation in circadian rhythms with the degree of sociality: interindividual variation increases with increased level of sociality. This study invites investigating the potential role of circadian rhythm variation in social organization.

